# Comparing Variant Call Files for Performance Benchmarking of Next-Generation Sequencing Variant Calling Pipelines

**DOI:** 10.1101/023754

**Authors:** John G. Cleary, Ross Braithwaite, Kurt Gaastra, Brian S. Hilbush, Stuart Inglis, Sean A. Irvine, Alan Jackson, Richard Littin, Mehul Rathod, David Ware, Justin M. Zook, Len Trigg, Francisco M. De La Vega

## Abstract

**Summary:** To evaluate and compare the performance of variant calling methods and their confidence scores, comparisons between a *test call* set and a *“gold standard”* need to be carried out. Unfortunately, these comparisons are not straightforward with the current Variant Call Files (VCF), which are the standard output of most variant calling algorithms for high-throughput sequencing data. Comparisons of VCFs are often confounded by the different representations of indels, MNPs, and combinations thereof with SNVs in complex regions of the genome, resulting in misleading results. A variant caller is inherently a classification method designed to score putative variants with confidence scores that could permit controlling the rate of false positives (FP) or false negatives (FN) for a given application. Receiver operator curves (ROC) and the area under the ROC (AUC) are efficient metrics to evaluate a test call set versus a gold standard. However, in the case of VCF data this also requires a special accounting to deal with discrepant representations. We developed a novel algorithm for comparing variant call sets that deals with complex call representation discrepancies and through a dynamic programing method that minimizes false positives and negatives globally across the entire call sets for accurate performance evaluation of VCFs.

**Availability:** RTG Tools is implemented as a multithreaded Java application and source code is available under BSD license at: https://github.com/RealTimeGenomics/rtg-tools

**Supplementary information:** Supplementary data are available at *Bioinformatics online*.

## 1 INTRODUCTION

High-throughput sequencing enables the identification of genetic variants in whole human genomes and exomes at a scale that is useful for population studies (1000 Genomes Project Consortium *et al.*, 2012) and clinical applications (Yang *et al.*, 2013). The output of such process is typically a set of variant, or indeed genotype “calls” (for diploid organisms) in the form of a Variant Call File (VCF), following the definitions set by the community standard (Danecek *et al.*, 2011).

When developing and validating sequencing pipelines and variant calling algorithms, the comparison of variant call sets is a common aim. One problem for benchmarking pipelines has been the lack of gold standards to which comparisons should be made and instead researchers have resorted to drawing Venn diagrams, which cannot resolve what caller works better (ORawe *et al.*, 2013). With the advent of comprehensive gold standards for selected human samples (Zook et al., 2014) the latter problem is going away, but the question of how to appropriately compare VCFs remains.

The naϊve way of comparing VCFs is to look at the same reference genome locations in the *baseline* (i.e. the gold standard) and *called* (test) variant call sets, and see if variants and genotype calls match at the same position. However, complications arise due to possible differences in representation for indels and complex variants between the baseline and the call sets. This often happen for indels within repeats or homopolymers, where their positions can be ambiguous due to alignment artifacts and how the variants are aligned with respect to the 5′ or 3′ ends of the reference. Another problem arises with multiple-nucleotide polymorphisms (MNPs) and complex variants which encompass combinations of simpler variants where calls can be locally phased (see Supplementary Note for examples).

`VCFtools` is a commonly used tool to manipulate and compare VCF files (Danecek *et al*., 2011). However, its comparison function does not deal with the complex call representations issues and it is slow. Other tools have appeared recently that deal with some of these problems, such as `SmaSH` (Talwalkar *et al*., 2014), `VCFlib` and `bcbio.variation`, typically by reducing complex calls into “primitives” (i.e. decomposing complex calls and MNPs into individual SNVs). Besides primitive representations being erroneous in some cases (a MNPs is a linked set of nucleotides, not independent SNVs), these methods still suffer from confounding given the multiple haplotypes possible, and do not attempt to globally optimize the comparison between call set and baseline in any way, resulting in a larger number of discrepancies than warranted between the two.

Here we present `vcfeval`, an algorithm that is implemented as part of the freely available `RTG Tools` software package that include functions for the fast manipulation and analysis VCF files, correctly dealing with variant representation confounders, optimizing globally to minimize discrepancies between a call set and the baseline, and providing utilities to perform ROC curve analysis, filtering and annotation of variant calls. We demonstrate that our method returns more accurate comparisons than previous methods and showcase a number of examples where others methods stumble but `vcfeval` returns correct results.

## 2 METHODS

### 2.1 Assumptions

`vcfeval` aims to match a baseline and called variants so as to maximize true positives (TPs) and minimize false positives (FPs) and false negatives (FNs) across the entire variant call set, which could include calls for an entire human genome. This is done in such a way that the number of TPs plus the number of FNs equals the total number of calls in the baseline for proper accounting and generations of receiver-operator curves (ROC).

Three pieces of information are needed when evaluating call sets of a reference sample: i) the reference sequence against which the variants were called (which needs to be the same; for example, a given reference genome assembly); ii) a baseline set of variants on the reference sample (the ground truth); and iii) a called set of variants on the reference sample. The called variant set will be the best possible if it correctly includes everything in the baseline (TPs), has no incorrect calls (FPs), and has not missed any call in the baseline (FNs). To deal with variant representation confounding, `vcfeval` implements a method that *replays* the variants from both the baseline and called sets to the reference genome assembly in a uniform way. While this normalization simplifies comparisons, replay does not guarantee a unique set of TP, FP, and FNs, as there might be alternative representations (haplotypes) at a given locus. Therefore, a global optimization method to select the most parsimonious among these options is needed. To simplify the exposition of the algorithm, we will first present the case of comparing two haploid call sets before proceeding to the more common diploid case.

### 2.2 Haploid Case

Let *R* = {*r*_1_,*r*_2_,…,*r*_*m*_} denote a reference nucleotide sequence of length *m*. Represent a haploid variant *v* on *R* by a triple (*a,s,e*) where *a* is the nucleotide sequence (possibly of length 0) of the variant allele, s ≥ 1 is the inclusive start locus and e ≤ *m* + 1 is the exclusive end locus. Given a variant *v* = (*a,s,e*), we define accessor functions *a(v) =a, s(v) = s*, and *e(v) = v*. The cardinality of a sequence or vector *X* is denoted by |*X*|.

Given a vector of variants *V* = (*v*_1_,*v*_2_, …,*v*_*n*_) with *s*(*v*_*k*_) ≥*e*(*v*_*k-1*_), we recursively define a haplotype function, *h*, that replays the variants *V* into the reference *R* as follows:

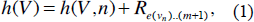

where

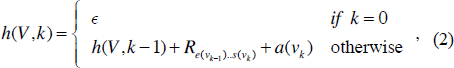

where + is understood to mean concatenation, ∊ denotes the empty string, and *R*_*α, β*_ indicates the reference nucleotides from position α up to but excluding β. Any nucleotide within *h*(*V*) is either drawn from the reference or from a variant in *V*.

Let *B* denote the set of baseline variants and *C* denote the set of called variants. We seek sequences of variants *X* and *Y* which maximize the number of agreements in the set *X*:

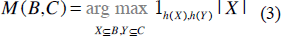

where the maximization is over all possible subsequences of variants *X* in *B* and variants *Y* in *C* and 1_*h(X),h(Y)*_ is the indicator function that is 1 iff the nucleotide sequences *h*(*X*) and *h*(*Y*) are the same; that is, *h*(*X*) = *h*(*Y*).

### 2.3 Diploid Case

Generalizing to the diploid case, each variant now has two alleles (not necessarily distinct) which we represent by (*a*_0_: *a*_1_, *s*, *e*). We introduce a phasing vector *P*_*v*_ =(*p*_1_,*p*_2_,*…p*_*n*_) **∊**{0,1}^*n*^, where *P*_*i*_, is 0 or 1 for each variant in *X* indicating whether *a*_0_ or *a*_1_ is used, respectively. Given a phasing vector *P*_v_, we use *P*_*v′*_, to denote the alternate phasing vector: defined by, *P*_*v′*_ =(l–*p*_1_,l–*p*_2_,.. l–*p*_*n*_). Thus if *P*_*v*_ represents the alleles selected by one of the haplotypes through the variants, then *P*_*v****′***_, represents the alleles in the alternate haplotype.

The haplotype function is modified to take the phasing vector into account:

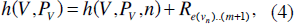

Where

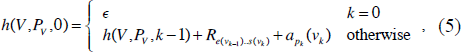

The corresponding global maximization is now additionally over all possible phasings of the variants:

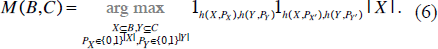

### 2.4 Path Creation

A *path* through a variant sequence is a selection of a subset of variants along with a choice in their phasing. `vcfeval` selects baseline and called *paths* ensuring that the variants included in these will be equal (after being replayed): these are thus classified as TPs. The calls excluded from the baseline *path* will correspond to FNs and the calls excluded from the called variant *path* will be classified as FPs. As noted above, potentially there are an exponential number of cases to be explored. Given diploid variant sets *B* and *C*, there are 3^|*B*|+|*C*|^ possible combinations of *X*, *P*_*X*_, *Y*, and *P*_*Y*_, since for each variant, *i*, there is the possibility for it to be excluded, included with *p*_*i*_ = 0, or included with *p*_*i*_ = 1.

We call a partially constructed *X* and *P*_*X*_ a *half path*. So a *half-path* through a set of variants is a selection of a subset of variants along with a choice in their phasing. Two half-paths taken together, *i.e*. a combination of *X*, *P*_*x*_, *Y*, *P*_*y*_, is a *path*.

It is possible to ‘step’ through each path, generating the full sequence of nucleotides in the (diploid or haploid) haplotypes represented by the *path*. This is termed *replaying* the path, resulting in a partial version, *h_α..β_*, of the *h* function in the formulation above. Further, for a position *r* in a reference of length *m*, provided 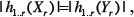 the indicator function can be factored as

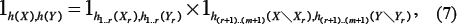

where 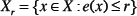 is those variants of *X* corresponding to the reference region up to and including *r*.

In practice, the number of paths to be explored can be reduced in three ways:

a. For homozygous variants both phasings are effectively the same, so only one of them need be explored.
b. Usually it is possible to determine that the indicator function will be 0 without computing the entire path. Once the indicator function is determined to be 0, the path can be discarded.
c. Finally, if two paths have reached the same reference position, *r*, and have the same replayed haplotype lengths, we retain the one maximizing **|** X**|** since any extension of these paths is independent of the path so far.

More precisely, for case (*c*) in a haploid situation, let *S = (X*_*r*_,*Y*_*r*_) and *S′* = (*X*_*r″*_,*Y*_*r″*_) be two partial paths such that *r* = *r*′ and 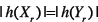 and 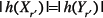. These conditions ensure *B*″ = *B*∖*B*_*r*_ and *C*″ = *C*∖*C*_*r*_ (that is, the remaining candidate variants) are the same for both paths and that we can split the indicator function using Eq. No. 7. The contribution to |X| in the maximal solution of the remaining portion (*r*+ l).. (*m* + l) will then be the same for both paths:

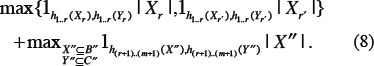

Since the maximization over *B*″ and *C*″ is the same for all extensions of *S* and *S*′, the global optimum solution for such a pair is determined by the optimal of the initial portion. The existence of the recurrence (8) permits a dynamic programming solution to be applied. With the addition of a phasing vector and dual haplotypes as per (3), these same simplifications to the computation can be applied in a diploid situation. Figure 1 shows a representation of best paths for a set of variants in the baseline and called sets (see the Supplementary Note for additional examples).

**Figure 1.**
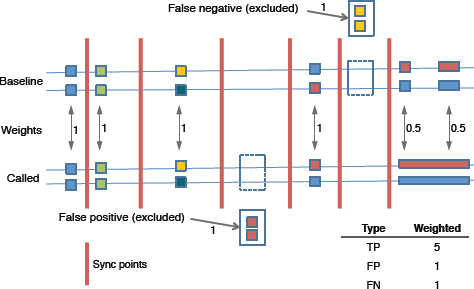
Best paths for baseline and called variants showing cases for false positives and negatives. Vertical lines are sync points. The inset table shows the total counts for the example after weighting.

### 2.5 Dynamic Programming Implementation

`vcfeval` selects baseline and called paths ensuring that the variants included in these will be equal (after being replayed); these included variants are classified as TPs. The calls excluded from the baseline path will correspond to FNs and the calls excluded from the called variant path are classified as FPs. (This notion of ‘correctness’ is with respect to the baseline, one could equivalently think of `vcfeval` as performing an advanced set intersection between two variant sets, and the terminology could alternatively be expressed as *B* ⋂ *C*, *B*∖*C*, *C*∖*B*, respectively.)

`vcfeval` takes a dynamic programming approach to finding the best path of replayed variants that globally maximizes TPs and minimizes FPs and FNs, by incrementally stepping through the haplotypes of each of the paths in a current working set of candidate paths. The size of the current working set is kept as small as possible by using the simplifications detailed above.

At any point during path replay, the next nucleotide to be generated will either be a reference base (if the haplotype is not currently within a variant), or a base from a variant allele (there is no particular distinction between a reference versus alternate allele). A path can be incrementally constructed during replay by delaying the decision of whether to exclude or include each variant until the replay has reached a point on the reference where that decision must be made in order to output the next haplotype base. At that point, the path is forked into three alternatives: one excluding the variant; one including the variant in a default phase; and one including the variant in the opposite phasing (actually by case (*a*) this last alternative is only required for heterozygous variants). Each of the alternatives is added to the working set, replacing the original.

Any path that results in haplotype disagreement during replay can be discarded from the working set according to case (*b*). If two (replayed) paths converge at the same position on the reference such that case (*c*) can be applied, then a path which maximizes the number of true positives up to that point can be kept and the other discarded. Partial paths which have converged, are said to be *c-equivalent.*

In practice such situations happen frequently keeping the memory and processing requirements reasonable. Note that our method considers zygosity when comparing variants (diploidy is assumed, except for sex chromosomes in humans), and thus to match, variants should have the same genotype (for some applications this requirement can be relaxed to allow a more lenient comparison that does not penalize mis-calling heterozygous as homozygous and vice versa). See the Supplementary Note for the pseu-do-code of the path creation and dynamic programing algorithms.

All heterozygous variants are treated as non-phased, allowing the best path procedure to discover the consistent phasing. Indeed, the phasing vector *P*_*v*_ in the best solution permits comparison with a provided phasing to measure phasing consistency, count switch errors, and so on. In principle, this algorithm could incorporate phasing information when provided in the input call-sets, by only including variants in a path having phasing consistent with the phase *P*_*v*_ of already added variants in the same phase set, thereby further reducing the search space. However, the current implementation is more general and performs well in practice.

### 2.6 Weighting

When comparing different variant call sets the number of true positives plus the number of false negatives should equal the total number of calls in the baseline, rather than being dependent on representational conventions of a particular call set. Generally, each called variant will have a corresponding baseline variant, but due to both variant representation confounding and repeat structures in the genome, there can be a many-to-many relationship between baseline and called mutation (see examples in the Supplementary Note). To keep the number of TPs plus the number of FNs equal to the total number of variants in the baseline, each called TP call must be weighted. To avoid pitfalls due to ambiguity when looking for equivalences in repeat regions, we perform weighting within unambiguous boundaries in the path. A *sync point* is a location where baseline and called paths are at the same position on the reference and they are not currently in the middle of any variant location after replay (see Figure 1 and Supplementary Note). An optimization in the *best path* creation skips all the genomic locations which does not contain any variants, thus the sync points occurs just before the next available variant. Once all the sync points are created, each called variant is weighted using following formula

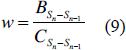

where *B* is the number of baseline variants between the current (*S*_*i*_) and previous sync points (*S*_i-1_) and *C* is the number of called variants between the current and previous sync points. Figure 1 shows an example of the results of weighting when a complex call is represented as a single variant in the call set, but as two variants in the baseline. The inset table in Figure 1 shows the total numbers of TP/FP/FN for this case after weighting. `vcfeval` outputs the sets of TP, FP and FN as separate VCF files for further analysis (see additional examples in the Supplementary Note).

### 2.7 Identifying variants of discrepant representation

To find complex variant examples to test variant comparison tools, we selected calls from two high-confidence variant call sets for the human cell line NA12878 which is the pilot reference material of the Genome-in-a-Bottle (GiaB) consortium: a) the NIST arbitration v.2.19 high-confidence calls (Zook *et al*., 2014) (hereafter ARB), and b) the RTG phasing consistent calls (Cleary *et al.,* 2014) (hereafter PF1S). These two VCFs files were chosen because they prefer to output complex variants in different ways. ARB tends to separate complex events into simple SNPs and indels without local phasing information whereas PHS tends to output complex variants as block substitutions and includes pedigree-based phasing for variants.

To find these complex regions, first we used `BedTools` v.2.22.1 to select variants from the ARB calls that were in regions in which there was **<** 50 bp between variants. Then, we added 49 bp on either side of these regions and then merged regions within 50 bp of each other to make sure we capture different representations of complex variants. Then, to distinguish between regions that had only SNPs from regions with at least one indel, we annotated each region with the number of SNPs and indels from ARB (see Supplementary Materials). Note that complex variants in ARB are in the default output format of GATK HaplotypeCaller v.2.8, which prefers to output complex events as separate SNPs and indels rather than as block substitutions. Next, we used `BedTools` to select variants in ARB and PHS VCFs that were inside the complex variants bed files calculated above. Finally, we used `vcfeval` v.3.4.2 to compare these two VCFs containing different representations of complex variants. We manually inspected sites that were deemed discordant and found that they were in fact discordant, either because the calls were completely missing in one of the VCFs or because only part of a complex event called by PHS was in ARB. For these partially called sites (where part of a complex event is called in one file and the full event is called in the other file), some normalization/comparison tools will find that one part is the same while the other is different.

### 2.8 Comparison of vcfeval with other tools

For normalization of variant representations, we tested: a) norm from `bcftools` v.7fa0d25 downloaded from GitHub on 2/19/15; b) `vt` v.b8219fd downloaded from GitHub on 3/5/15; c) `SMaSH v.0ff627a` downloaded from GitFlub on 3/5/15; d) `vcfallel-icprimitives` from `vcflib` downloaded from GitHub on 3/6/15; e) `bcbio.variation` v.0.2.4 prep and normalize.

For variant comparison, we tested: a) `vcfintersect` from `vcflib` downloaded from GitFlub on 3/6/15, before and after normalization with `vcfallelicprimitives`; b) `stats` from `bcftools` v.7fa0d25 downloaded from GitFlub on 2/19/15, before and after normalization with `bcftools` norm; c) diff from `vcftools` v.0.1.12b after normalization with vt or `bcftools` norm; d) *smash* python v.0ff627a downloaded from GitFlub on 3/5/15 with -- `normalization` flag; e) smash `calldif` f v.0ff627a downloaded from GitFlub on 3/5/15 without normalization; f) `bcbio. variation` v.0.2.4 variant-compare with `bcbio.variation prep` and `normalize`

### 2.9 Generation of ROC curves

`vcfeval` provides a number of useful outputs to draw ROC curves and for the debugging of variant calling algorithms, including VCFs with the TP, FN, and FP of the test call set. The `rocplot` command of `RTG Tools` allows the user to select one or multiple comparisons to the same baseline, select the score to use for sorting the points, and in interactive mode provides the number of TP, precision, recall (sensitivity), and F-measure at a given threshold score in a plot. For the ROC curves presented in Figure 2, we compared call sets produced by a number of callers with the PHS baseline. The input for the variant calling pipelines was FastQ data for run ERR174327 available in the EBI ENA in project Acc. No. PRJEB3246. We selected a subset of lanes to reach about 40X depth of 2xl00bp paired reads for the sample NA12878 produced on the HiSeq 2000 by Illumina (<u>http://www.illumina.com/platinumgenomes/</u>). This data was mapped to the hgl9 human reference genomes with decoys (1000 Genomes Project Consortium *et al.*, 2012) and variants were called by the following pipelines: i) BWA-MEM v0.7.10/GATK UG v3.2.2 (99.5% SNV and indel 99% cut-off%); and ii) BWA-MEM V0.7.10/GATK HC v 3.2.2 (99.5% SNV and indel 99% cut-ofP/o); and iii) RTG Variant 3.3.2 (RTG map and snp).

**Figure 2.**
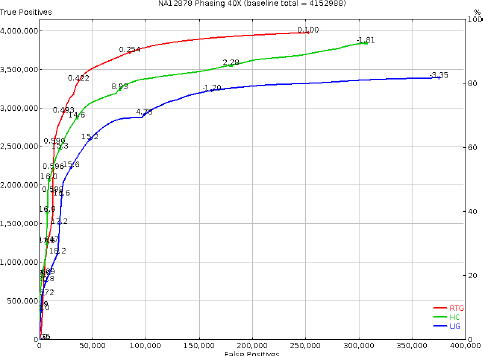
ROC curve comparing multiple call sets from the Illumina Platinum data for NA12878 (2xl00bp, 40X) made with GATK Unified Geno-typer (UG) and Haplotype Caller (HC) v3.2 (DePristo *et al*, 2011) and RTG Variant v3.3 `(RTG) versus` the PHS gold standard (Cleary *et al.*, 2014), sorted by recalibration scores (VQSLOD for GATK and AVR for RTG, some thresholds indicated in the lines).

### 2.10 Other utilities of RTG Tools

In addition to `vcfeval`, the `RTG Tools` software package includes a number of utility functions to manipulate and analyze VCFs, some of which are summarized here. The command `vcf-stats` provides summary statistics for the VCFs, such as the number of each type of variants and global quality metrics like Ti/Tv and the heterozygote/homozygote ratios. When variants are called in related samples a statistic that correlates with the quality of the call set is the number of Mendelian inconsistency errors (MIEs). The `mendelian` command performs this count when provided with a PED file describing the family structure (see Supplementary Note for output example). Other common tasks supported include the merging (`vcfmerge`), splitting (`vcf subset`), annotation (`vcfannotate`), creating tabix index (`index`), and filtering of VCF files by scores or annotations (`vcffilter`). Note that most `RTG Tools` commands require that the VCF tabix index is available and that the reference used in variant calling is present in an internal format - the SDF format (Cleary *et al.,* 2014) - which facilitates indexing and storage of associated metadata such as chromosome ploidy, PAR region location, etc. Thus, we provide the command `format` to create an SDF version of the reference from FASTA files. Further, `RTG Tools` can operate in block compressed VCFs directly to save space, and provides a command to block-compress such files (`bgzip`).

## 3 RESULTS

### 3.1 Comparative performance of vcfeval

When variants are near each other and within repetitive regions there are several possible representations of them in the VCF format. To find complex variant examples to test comparison tools, we identified regions containing variants within 50bp of each other in the arbitration calls developed by Zook *et al.* (2014) for the sample NA12878 (ARB), and compared these to phasing consistent calls we derived from the analysis of the 17 member CEPH 1463 pedigree to which the sample belongs (PHS) (Cleary *et al*.,2014).

In the regions containing only SNPs, we found that 546,712 calls from the ARB VCF were consistent with 526,928 calls in the PHS VCF. Of these calls, 41,572 have different representations (based on them being determined to be discordant when using `bcftools stats` without normalization). In the regions containing one or more indels, we found that 184,634 calls from the ARB VCF were consistent with 148,833 calls in the PHS VCF. Of these calls, at least 70,239 have different representations (based on them being determined to be discordant when using `bcftools stats` without normalization). Note that the remaining calls may have the same representation apart from differences in phasing information.

We next tested whether other normalization and comparison approaches are able to determine that the calls are consistent when `vcfeval` determines they are consistent. From each comparison tool, we tallied sites the tool determined to be specific to ARB VCF, sites specific to PHS VCF, sites in ARB that were also in PHS (Both), and sites that the tool could not compare because they were too complex or overlapping (Table 1).

**Table 1.**
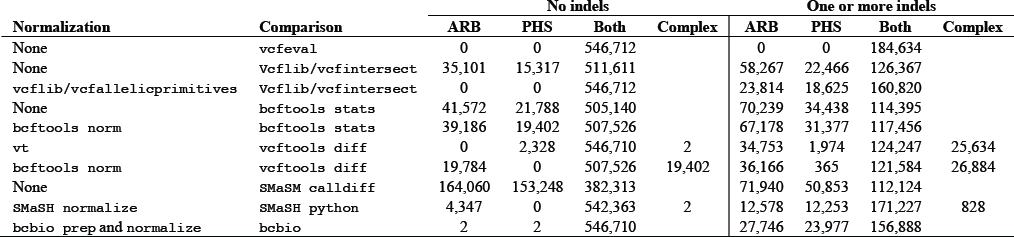
Comparison of normalization and VCF comparison tools. We provided as input two VCFs (ARB, NIST arbitration or PHS, RTG Phasing), either as raw calls or in pre-normalized format, to a number of comparison tools and counted the number of variants that were deemed discrepant for each of the files. Since some tools deal differently with SNVs vs indels, we performed the test with VCFs where all indels were filtered, or with the full set of calls including indels. The column “Complex” lists sites that were deemed too complex for the tool to assess.

| Normalization | Comparison | No indels |  |  |  | One or more indels |  |  |  |
| --- | --- | --- | --- | --- | --- | --- | --- | --- | --- |
|  |  | ARB | PHS | Both | Complex | ARB | PHS | Both | Complex |
| None | `vcfeval` | 0 | 0 | 546,712 |  | 0 | 0 | 184,634 |  |
| None | `VcfLib/vcfintersect` | 35,101 | 15,317 | 511,611 |  | 58,267 | 22,466 | 126,367 |  |
| `vcflib/vcfallelicprimitives` | `VcfLib/vcfintersect` | 0 | 0 | 546,712 |  | 23,814 | 18,625 | 160,820 |  |
| None | `bcftools stats` | 41,572 | 21,788 | 505,140 |  | 70,239 | 34,438 | 114,395 |  |
| `bcftools norm` | `bcftools stats` | 39,186 | 19,402 | 507,526 |  | 67,178 | 31,377 | 117,456 |  |
| `vt` | `vcftools diff` | 0 | 2,328 | 546,710 | 2 | 34,753 | 1,974 | 124,247 | 25,634 |
| `bcftools norm` | `vcftools diff` | 19,784 | 0 | 507,526 | 19,402 | 36,166 | 365 | 121,584 | 26,884 |
| None | `SMAsh calldiff` | 164,060 | 153,248 | 382,313 |  | 71,940 | 50,853 | 112,124 |  |
| `SMAsh normalize` | `SMAsh python` | 4,347 | 0 | 542,363 | 2 | 12,578 | 12,253 | 171,227 | 828 |
| `bcbio prep and normalize` | `bcbio` | 2 | 2 | 546,710 |  | 27,746 | 23,977 | 156,888 |  |

We found `vcflib vcfintersect` with `vcfallel-icprimitives` was the only other tool able to determine that all of the sites were consistent in the VCF files without indels, though `bcbio.variation` incorrectly called 2 sites inconsistent out of 546,712. No tool except for `vcfeval` was able to determine that all of the sites in the regions containing indels were consistent. The python version of `SMaSH` came closest, determining that 171,227 of 184,634 sites were consistent, but it was not able to handle compound heterozygous variants. Even though the Java version of `SMaSH (calldiff)` has a phasing-aware realignment algorithm, it determined that the fewest number of sites were consistent. This low number is because it requires that both VCF files have exactly the same phasing information, and ABR VCF does not contain phasing information while the PHS VCF does.

### 3.2 ROC curves

Once the variant representation problem has been addressed at the global level for a call set, comparison of test VCFs to a gold standard can be performed. Since NGS technology is not perfect and errors need to be distinguished from sequencing errors, variant calls usually come accompanied with quality scores that try to provide an assessment of the probability a new allele (QUAL) or genotype call (GQ) is an error. Sometimes empirical recalibration scores based on machine learning approaches are preferred since is difficult to model all experimental artifacts in the NGS protocol (DePristo *et al*., 2011). Identifying FPs and FNs as compared with the gold standard for a given a number of sorted score thresholds permit the construction of receiver-operator curves (ROC) which can be used to assess the performance of such scores and to compare the performance of different variant calling pipelines. In a ROC curve the true positives and false positives are plotted over various threshold values. The thresholds commonly used in variant analysis are the genotype quality (GQ) or recalibration scores from the VCF file.

Figure 2 shows a ROC curve produced with the `rocplot` command from the output of `vcfeval` comparing several call sets with the PHS call set as baseline. A variant call set and scoring system will be considered best when the curve gets closer to the top right of the plot (i.e. larger AUC). Although TP and TN rates and other metrics can be calculated for a given score threshold in the curves, and these can be used for overall comparisons of sensitivity and specificity, the analysis of the ROC permits to better justify a given threshold and assess the discriminative power of the variant quality scoring or recalibration schemes of each method.

## 4 CONCLUSIONS

The genomics and medical genetics communities are rallying to develop reference standard samples and associated ground truth sets of variants to evaluate both experimental and bioinformatic pipelines for human whole-genome and exome sequencing production (Zook *et al*., 2014). The US National Institute of Standards (NIST), had recently released the first standard reference material for such applications (RM 8398 - Human DNA for Whole-Genome Variant Assessment - for the sample NA128768) and through the GiaB consortium gold standard variant sets have been generated in VCF format from multiple approaches to be used as ground truth in comparisons.

The availability of such references allows to objectively compare variant calling algorithms, identify their flaws, and drive algorithm improvement. This permits going beyond Venn diagrams (ORawe *et al*., 2013) and instead moving to ROC curve analysis (Cleary *et al*., 2014), much more useful in the assessment of high-throughput sequencing pipelines for clinical applications. This workflow requires the analysis of VCF files and the ability to accurately compare different VCFs with such baselines, appropriately dealing with complex variant representation issues and looking at the call sets as a whole.

We developed `vcfeval` to enable performing such comparisons in an optimized fashion together with a set of useful VCF manipulation tools that are fast, and easy to use. We show that `vcfeval` produces the most accurate and parsimonious comparisons between VCF files among existing tools and permits the analysis of ROC curves to select the optimal variant score threshold to achieve the TP/FP balance that a given application requires. If the downstream analytics could utilize the variant confidence scores rather than consider all provided variants as true, no filtering would be necessary, but the ROC analysis would still allow optimizing scoring and recalibration systems. Together with the additional utilities included in the `RTG Tools` package, we believe these tools will find extensive use. While our focus has been to compare a test call set with a baseline, which is assumed to be the ground truth, the algorithms we developed could be extended for the case of comparing multiple VCF files. Such is the case of the harmonization of calls coming from single sample pipelines or different sequencing platforms, a topic which will be the subject of future work.

The Global Alliance for Genomics and Health benchmarking team is currently working to develop standardized definitions for performance metrics (e.g., true positive, false positive, and false negative) to ensure comparability between benchmarking tools. The authors are working with this team to ensure `vcfeval` provides outputs consistent with these definitions

## ACKNOWLEDGEMENTS

We are grateful for the support of Graham Gaylard, Jason Blue-Smith, Stephen Lombardi (RTG), and Marc Salit (NIST). We also thank Tal Shmaya (Annai Systems) for providing an updated ROC figure. We dedicate this work to the memory of John G. Cleary. Certain commercial equipment, instruments, or materials are identified in this paper only to specify the experimental procedure adequately. Such identification is not intended to imply recommendation or endorsement by the NIST, nor is it intended to imply that the materials or equipment identified are necessarily the best available for the purpose.

*Conflict of Interest*: All authors, except for JMZ, were employed or received payments from Real Time Genomics, Inc. or its subsidiaries during the performance of this work.

## REFERENCES

1000 Genomes Project Consortium et al. (2012) An integrated map of genetic variation from 1,092 human genomes. Nature, 491, 56–65.

Cleary,J.G. et al. (2014) Joint variant and de novo mutation identification on pedigrees from high-throughput sequencing data. bioRxiv.

Danecek,P. et al. (2011) The variant call format and VCFtools. Bioinformatics, 27, 2156–2158.

DePristo,M. A. et al. (2011) A framework for variation discovery and genotyping using next-generation DNA sequencing data. Nat. Genet., 43, 491–498.

O’Rawe,J. et al. (2013) Low concordance of multiple variant-calling pipelines: practical implications for exome and genome sequencing. Genome Medicine, 5, 28.

Talwalkar,A. et al. (2014) SMaSH: a benchmarking toolkit for human genome variant calling. Bioinformatics.

Yang,Y. et al. (2013) Clinical Whole-Exome Sequencing for the Diagnosis of Mendelian Disorders. N Engl J Med, 131002140031007.

Zook,J.M. et al. (2014) integrating human sequence data sets provides a resource of benchmark snP and indel genotype calls. Nature Biotechnology, 1–8.

